# Distinct and sequential re-replication barriers ensure precise genome duplication

**DOI:** 10.1101/366666

**Authors:** Yizhuo Zhou, Pedro N. Pozo, Seeun Oh, Haley M. Stone, Jeanette Gowen Cook

## Abstract

Achieving complete and precise genome duplication requires that each genomic segment be replicated only once per cell division cycle. Protecting large eukaryotic genomes from re-replication requires an overlapping set of molecular mechanisms that prevent the first DNA replication step, the DNA loading of MCM helicase complexes to license replication origins. Previous reports have defined many such origin licensing inhibition mechanisms, but the temporal relationships among them are not clear, particularly with respect to preventing re-replication in G2 and M phases. Using a combination of mutagenesis, biochemistry, and single cell analyses in human cells, we define a new mechanism that prevents re-replication through hyperphosphorylation of the essential MCM loading protein, Cdt1. We demonstrate that Cyclin A/CDK1 hyperphosphorylates Cdt1 to inhibit MCM re-loading in G2 phase. The mechanism of inhibition is to block Cdt1 binding to MCM independently of other known Cdt1 inactivation mechanisms such as Cdt1 degradation during S phase or Geminin binding. Moreover, we provide evidence that protein phosphatase 1-dependent Cdt1 dephosphorylation at the mitosis-to-G1 phase transition re-activates Cdt1. We propose that multiple distinct, non-redundant licensing inhibition mechanisms act in a series of sequential relays through each cell cycle phase to ensure precise genome duplication.

**Author Summary:** The initial step of DNA replication is loading the DNA helicase, MCM, onto DNA during the first phase of the cell division cycle. If MCM loading occurs inappropriately onto DNA that has already been replicated, then cells risk DNA re-replication, a source of endogenous DNA damage and genome instability. How mammalian cells prevent any sections of their very large genomes from re-replicating is still not fully understood. We found that the Cdt1 protein, one of the critical MCM loading factors, is inhibited specifically in late cell cycle stages through a mechanism involving protein phosphorylation. This phosphorylation prevents Cdt1 from binding MCM; when Cdt1 can’t be phosphorylated MCM is inappropriately re-loaded onto DNA and cells are prone to re-replication. When cells divide and transition into G1 phase, Cdt1 is then dephosphorylated to re-activate it for MCM loading. Based on these findings we assert that the different mechanisms that cooperate to avoid re-replication are not redundant, but rather distinct mechanisms are dominant in different cell cycle phases. These findings have implications for understanding how genomes are duplicated precisely once per cell cycle and shed light on how that process is perturbed by changes in Cdt1 levels or phosphorylation activity.

## Introduction

During normal cell proliferation DNA replication must be completed precisely once per cell cycle. A prerequisite for DNA replication in eukaryotic cells is the DNA loading of the core of the replicative helicase, the <u>m</u>ini<u>c</u>hromosome <u>m</u>aintenance complex (MCM). The process of MCM loading is known as DNA replication origin licensing, and it is normally restricted to the G1 cell cycle phase [1–3]. In proliferating mammalian cells, hundreds of thousands of replication origins are licensed in G1, then a subset of these origins initiate replication in S phase. To achieve precise genome duplication, no origin should initiate more than once per cell cycle, and preventing re-initiation is achieved by preventing re-licensing [4–7]. Improper re-licensing in S, G2, or M phases leads to re-initiation and re-replication, a source of DNA damage and genome instability that can promote cell death or oncogenesis (reviewed in [7–10]).

Re-licensing is prevented by an extensive collection of mechanisms that inhibit the proteins required to load MCM. In vertebrates, multiple transcriptional and post-transcriptional mechanisms target each of the individual licensing components that load MCM complexes: the <u>o</u>rigin recognition <u>c</u>omplex (ORC), the Cdc6 (<u>c</u>ell <u>d</u>ivision <u>c</u>ycle 6), and Cdt1 (<u>C</u>dc10-<u>d</u>ependent transcript 1) proteins as well as MCM subunits themselves are all inactivated for licensing outside of G1 phase (reviewed in [1, 3, 7, 11–14]). These mechanisms include regulation of licensing components’ synthesis, subcellular localization, chromatin association, protein-protein interactions, and degradation. In addition, cell cycle-dependent changes in chromatin structure contribute to licensing control [15]. Why have mammals evolved so very many distinct molecular mechanisms to prevent re-replication? Are each of these mechanisms *redundant* with one another, or do they operate in a *temporal series* coupled to cell cycle progression? In this study we investigated potential differences between re-replication control *during* S phase and re-replication control *after* S phase ends. We considered that licensing control in late S phase and G2 phase is particularly important because the genome has been fully replicated by this time, and thus G2 cells have the highest amount of available DNA substrate for re-replication.

We were inspired to explore the notion of sequential re-replication control by studies of mammalian Cdt1. One of the well-known mechanisms to avoid re-replication in mammalian cells is degradation of Cdt1 during S phase. Beginning in late S phase however, Cdt1 re-accumulates and reaches levels during G2 phase similar to its levels in G1 phase when Cdt1 is fully active to promote MCM loading [16–21]. One mechanism to restrain Cdt1 activity in G2 is binding to a dedicated inhibitor protein, Geminin, which interferes with Cdt1-MCM binding [22–24]. Interestingly, mammalian Cdt1 is hyperphosphorylated in G2 phase relative to Cdt1 in G1 phase [16, 17], but the consequences of those phosphorylations are largely unknown. Here, we elucidated a novel phosphorylation-dependent mechanism that inhibits Cdt1 licensing activity in G2 and M phase rather than inducing Cdt1 degradation to ensure precise genome duplication. We propose that multiple re-licensing inhibition mechanisms are not redundant, but rather act in a sequential relay from early S phase (replication-coupled destruction) through mid-S phase (degradation plus geminin) to G2 and M phase (geminin plus Cdt1 hyperphosphorylation) to achieve stringent protection from re-replication for mammalian genomes.

## Results

### Cdt1 phosphorylation inhibits DNA re-replication

Mammalian Cdt1 is phosphorylated in G2 phase and mitosis [17, 19, 20], and we hypothesized that this phosphorylation contributes to blocking re-replication by directly inhibiting Cdt1 licensing activity. To test that hypothesis, we generated mutations in candidate phosphorylation sites illustrated in Fig. 1A. We first compared the activity of normal Cdt1 (wild-type, WT) to a previously-described Cdt1 variant, “Cdt1-5A” bearing mutations at five phosphorylation sites. We had shown that this variant, “Cdt1-5A” (S391A, T402A, T406A, S411A, and S491A) is both unphosphorylatable *in vitro* by stress-induced MAP kinases and compromised for G2 hyperphosphorylation detected by gel mobility shift [17]. Four of the five sites are in a region of low sequence conservation and high-predicted intrinsic disorder [25](Fig. 1A and Supplementary Fig. S1). This “linker” region connects the two winged-helix domains of Cdt1 that have been characterized for MCM binding (C-terminal “C domain”) [26] or for binding to the inhibitor Geminin (middle “M domain”) [27]. Both domains are required for metazoan licensing activity [28–32]. We inserted cDNAs encoding either wild-type Cdt1 (Cdt1-WT) or Cdt1-5A into a single chromosomal FRT recombination site under doxycycline-inducible expression control in the U2OS cell line. All Cdt1 constructs bear C-terminal HA epitope and polyhistidine tags to distinguish ectopic Cdt1 from endogenous Cdt1.

**Figure 1.**
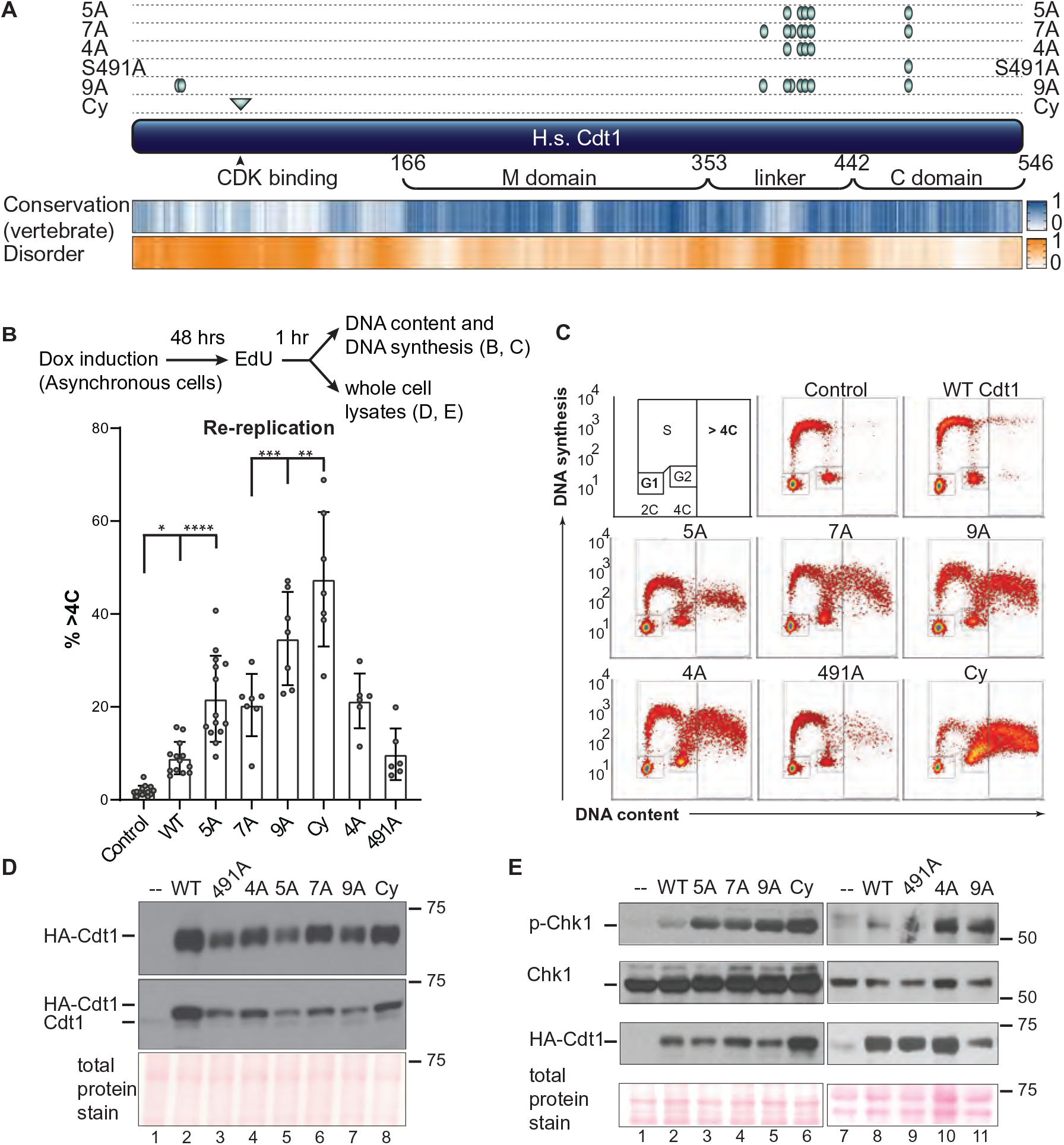
Cdt1 phosphorylation restrains re-replication. **A)** Schematic of the human Cdt1 protein illustrating features and variants relevant to this study. Cdt1 contains two structurally characterized domains, the Geminin and MCM binding domain (M) and a C-terminal MCM binding domain (C). The Ser/Thr-Pro sites that were altered for this study are marked with green ovals, and the cyclin binding motif is marked with a green triangle. Positions are T29, S31, S372, S391, S394, T402, T406, S411, and S491; the cyclin binding motif (Cy) is 68-70. Human Cdt1 was aligned with 26 vertebrate Cdt1 sequences using ClustalW, and a relative conservation score was derived (see also Methods and Supplementary Fig. S1). The blue heatmap indicates relative conservation at each amino acid position of human Cdt1. An intrinsic disorder score was also derived for human Cdt1 and shown as the corresponding orange heatmap. Darker shades indicate greater conservation or disorder respectively. **B)** Asynchronously growing U2OS cells with the indicated chromosomally-integrated inducible Cdt1 constructs were treated with 1 µg/mL doxycycline for 48 hours and labeled with EdU for 1 hour before harvesting. Cells were analyzed by flow cytometry for DNA content with DAPI and for DNA synthesis by EdU detection; the workflow is illustrated at the top. The bar graph plots the percentages of re-replicating cells across all experiments. Bars report mean and standard deviations. Asterisks indicate statistical significance determined by one-way ANOVA (*p=0.0175, **p=0.0023, ***p= 0.007, **** p<0.0001); 5A vs 7A, 5A vs 4A and WT vs 491A were not significant as defined by p>0.05. **C)** One representative of the multiple independent biological replicates summarized in B is shown. **D)** Whole cell lysates as in B were subjected to immunoblotting for ectopic (HA) or endogenous and ectopic Cdt1; Ponceau S staining of total protein serves as a loading control. **E)** Asynchronously growing U2OS cells were treated with 1 µg/mL doxycycline for 48 hours, and whole cell lysates were probed for phospho-Chk1 (S345), total Chk1, HA-Cdt1, and total protein; one example of at least two independent experiments is shown.

As a measure of relative Cdt1 activity, we induced Cdt1 production to approximately 5-10 times higher levels than endogenous Cdt1 in asynchronously proliferating cells over the course of 48 hrs (Fig. 1D, compare lanes 1 and 2). The amount of re-replication induced by Cdt1 overproduction is directly related to Cdt1 licensing activity [30]. As previously reported [33, 34], Cdt1-WT overproduction in human cells induced some re-replication, which we detected by analytical flow cytometry as a population of cells with DNA content greater than the normal G2 amount (>4C, Fig. 1B and 1C, and Supplementary Figure S2A). Strikingly however, overproducing Cdt1-5A (Fig. 1D, lane 5) induced substantially more re-replication suggesting that this variant is intrinsically more active (Fig 1B and 1C). DNA re-replication can also induce the formation of giant nuclei [35, 36], and we noted that the average nuclear area of cells overproducing Cdt1-WT was somewhat larger than control nuclei, whereas nuclei of cells overproducing Cdt1-5A were even larger (Supplementary Fig. S2A). Thus, Cdt1-5A expression not only induces more cells to re-replicate, but it also induces a higher degree of re-replication in those individual cells compared to Cdt1-WT expression.

Re-replication is an aberrant genotoxic phenomenon characterized by molecular markers of DNA damage (reviewed in [1, 7, 14]). As an additional measure of re-replication, we analyzed lysates of Cdt1-overproducing cells for Chk1 phosphorylation, a marker of the cellular DNA damage response. Cdt1-5A consistently induced more Chk1 phosphorylation than WT Cdt1 (Fig. 1E, compare lanes 2 and 3). Moreover, cells overproducing Cdt1-5A were also ∼3 times more likely to generate γ-H2AX foci, another marker of re-replication-associated DNA damage [37] (Supplementary Fig. S2B). We also noted that the accumulation of re-replicated cells came at the expense of G1 cells, consistent with a scenario in which re-replication during S or G2 induced a DNA damage response and a G2 checkpoint cell cycle arrest (Supplementary Fig. S3).

Phosphorylation at two additional candidate CDK/MAPK target sites in the linker region has been detected in global phosphoproteomics analyses [38]. To test the potential additional contribution of these sites to Cdt1 regulation, we included the mutations S372A and S394A to Cdt1-5A to create Cdt1-7A (Fig. 1A). Cdt1-7A overproduction did not induce more re-replication or DNA damage than Cdt1-5A (Fig. 1B and 1C, p>0.05, Fig. 1E, lane 4). From this observation, we infer that Cdt1-5A is already at the maximal deregulation that is achievable from phosphorylation in the linker region, and that additional phosphorylations do not further affect activity. (Of note, the y-axis values of re-replicating cells varies among different mutants and reflects snapshots of the rates of DNA synthesis in the final 30 minutes of Cdt1 expression.) To assess the importance of the four sites in the linker relative to the single site in the C-terminal domain, we generated Cdt1-4A and Cdt1-S491A (Fig. 1A). Cdt1-4A was as active as Cdt1-5A for inducing re-replication, whereas Cdt1-S491A only induced as much re-replication as Cdt1-WT (Fig. 1B and 1C). Like Cdt1-5A, Cdt1-4A induced substantially more DNA damage (phospho-Chk1) than Cdt1-WT (Fig. 1E, lanes 8 and 10). Thus, linker region phosphorylation inhibits Cdt1 activity.

Cdt1 is also phosphorylated at both T29 and S31 [19, 38] (see also Fig. 1A). CDK-dependent phosphorylation at T29 generates a binding site for the SCF^Skp2^ E3 ubiquitin ligase, which contributes to Cdt1 degradation during S phase [34, 39, 40]. The stress MAPK JNK (c-Jun N-terminal kinase) has also been reported to inhibit Cdt1 by phosphorylating T29 [41]. To determine if these N-terminal phosphorylations collaborate with linker region phosphorylations, we added the two mutations, T29A and S31A, to Cdt1-7A to generate Cdt1-9A. Cdt1-9A overproduction induced somewhat more re-replication than the three Cdt1 variants bearing only linker region mutations, Cdt1-4A, 5A, and 7A (Fig. 1B and 1C), and Cdt1-9A induced similar amounts of DNA damage checkpoint activation as these three linker variants (pChk1, Fig. 1E lanes 5 and 11). As an additional test, we included in our analysis a Cdt1 variant with a previously-characterized mutation in the cyclin binding motif, Cdt1-Cy (RRL to AAA at positions 66-68, Fig. 1A) [40]. We expect that this alteration compromises phosphorylation at most/all CDK-dependent phosphorylation sites. As we had noted in a previous study [28], Cdt1-Cy sometimes accumulated to higher levels than Cdt1-WT, particularly after longer induction times (e.g. Fig. 1E, lane 6), and this variant induced the highest amount of both re-replication and Chk1 phosphorylation (Fig. 1B,1C, and 1E). We presume that higher Cdt1-Cy stability contributes to enhanced re-replication activity, but this effect must be independent of phosphorylation at T29 and S31 since Cdt1-Cy is more stable and more active than both Cdt1-9A and a previously-tested Cdt1 mutant “2A” [28].

### Cdt1 phosphorylation prevents MCM re-loading in G2 cells

Re-replication requires that MCM be loaded back onto DNA that has already been duplicated followed by a second round of initiation. We sought to determine when during the cell cycle the mutations that de-regulate Cdt1 activity induce MCM-reloading. For this test, we used an analytical flow cytometry assay that detects only bound MCM because we extract soluble MCM with detergent prior to fixing and anti-MCM staining [42, 43]. We focused on two of the Cdt1 variants, Cdt1-4A because it represents a fully de-regulated linker, and Cdt1-Cy which induced the most re-replication after 48 hours of expression (Fig. 1). In asynchronously proliferating cultures, we induced Cdt1 for 24 hours which is slightly more than one full cell cycle in these U2OS cells to allow all cells to pass through each cell cycle phase. Particularly because Cdt1-Cy accumulates faster than Cdt1-WT over time, we analyzed expression shortly after doxycycline induction (Fig. 2A). At this early time point, all three forms of ectopic Cdt1 (WT, 4A, and Cy) were produced at similar amounts (Fig. 2B.). We then subjected these parallel cultures to analysis of DNA content to indicate cell cycle phase (x-axes) and MCM loading (y-axes) (Fig. 2C). In control cells, MCM is rapidly loaded in G1 and then progressively removed throughout S phase (illustrated in Fig. 2C). Overproducing normal Cdt1(“WT”) for 24 hours had only minimal impact on this pattern. In contrast to Cdt1-WT, both the Cdt1-4A and Cdt1-Cy variants induced a striking “spike” of MCM loading in cells with 4C DNA content (i.e. G2 phase) (Fig. 2C and 2D). Of note, we did not detect aberrant MCM loading in either G1 or S phase cells. We had previously established that linker phosphorylations do not impair Cdt1 degradation during S phase [17], so we interpret these results as MCM re-loading only after S phase is complete. Since these cells only overproduced Cdt1 for one cell cycle we also conclude that re-loading occurs within a single cell cycle.

**Figure 2.**
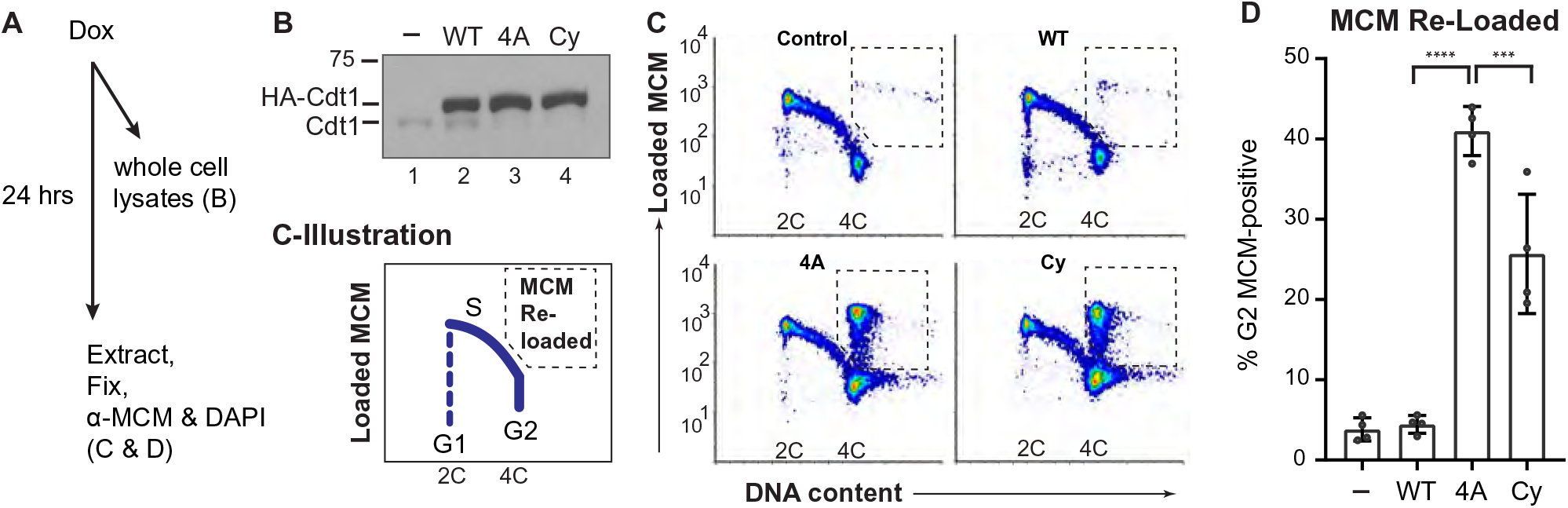
Cdt1 phosphorylation prevents MCM re-loading in G2 cells. **A)** Workflow: Asynchronously proliferating U2OS cells with inducible Cdt1 were treated with 0.05 µg/ml doxycycline then subjected to immunoblotting in B or analytical flow cytometry in C and D. **B)** Immunoblot analysis of initial Cdt1 expression 6 hrs after dox induction. Lysates were probed with anti-Cdt1 to detect both endogenous and ectopic Cdt1. **C)** Flow cytometry analysis of MCM loading 24 hrs after ectopic Cdt1 induction. Cells were detergent-extracted prior to fixation to remove unbound MCM, then stained for DNA content with DAPI (x-axes) and with anti-MCM2 as a marker of loaded MCM complexes (y-axes). One representative of multiple independent biological replicates is shown, and the illustration depicts typical positions of proliferating cells in G1, S, and G2 phase. The dashed boxes show the gates to quantify MCM re-loading in late S/G2 cells. **D)** Quantification of four independent replicates as in C. The bars report means and standard deviations. Asterisks indicate statistical significance determined by one-way ANOVA (***p= 0.0002, **** p<0.0001); Control vs WT was not significant as defined by p>0.05.

We had previously shown that normal Cdt1 from lysates of nocodazole-arrested (early mitotic) HeLa cells typically migrated slower than Cdt1-5A by standard SDS-PAGE [17]; we made similar observations in U2OS cells synchronized by S phase arrest then release into nocodazole (synchronization and expression strategy in Fig. 3A; Cdt1 migration by standard SDS-PAGE in Fig. 3B, middle panel lanes 2 and 5). As a more quantitative and consistent measure of Cdt1 phosphorylation, we analyzed Cdt1 migration in the presence of Phos-tag reagent which retards protein mobility proportional to the extent of phosphorylation [44]. HA-Cdt1 from nocodazole-arrested cells is a mixture of slow-migrating species on Phos-tag gels compared to HA-Cdt1 from G1 cells, and this migration was accelerated by phosphatase treatment of the lysates *in vitro* prior to electrophoresis (Supplementary Fig. S4A). The distribution of ectopic Cdt1-5A bands was lower than Cdt1-WT bands on Phos-tag gels (Fig. 3B, lanes 2 and 5), demonstrating that these sites are among the sites phosphorylated late in the cell cycle.

**Figure 3.**
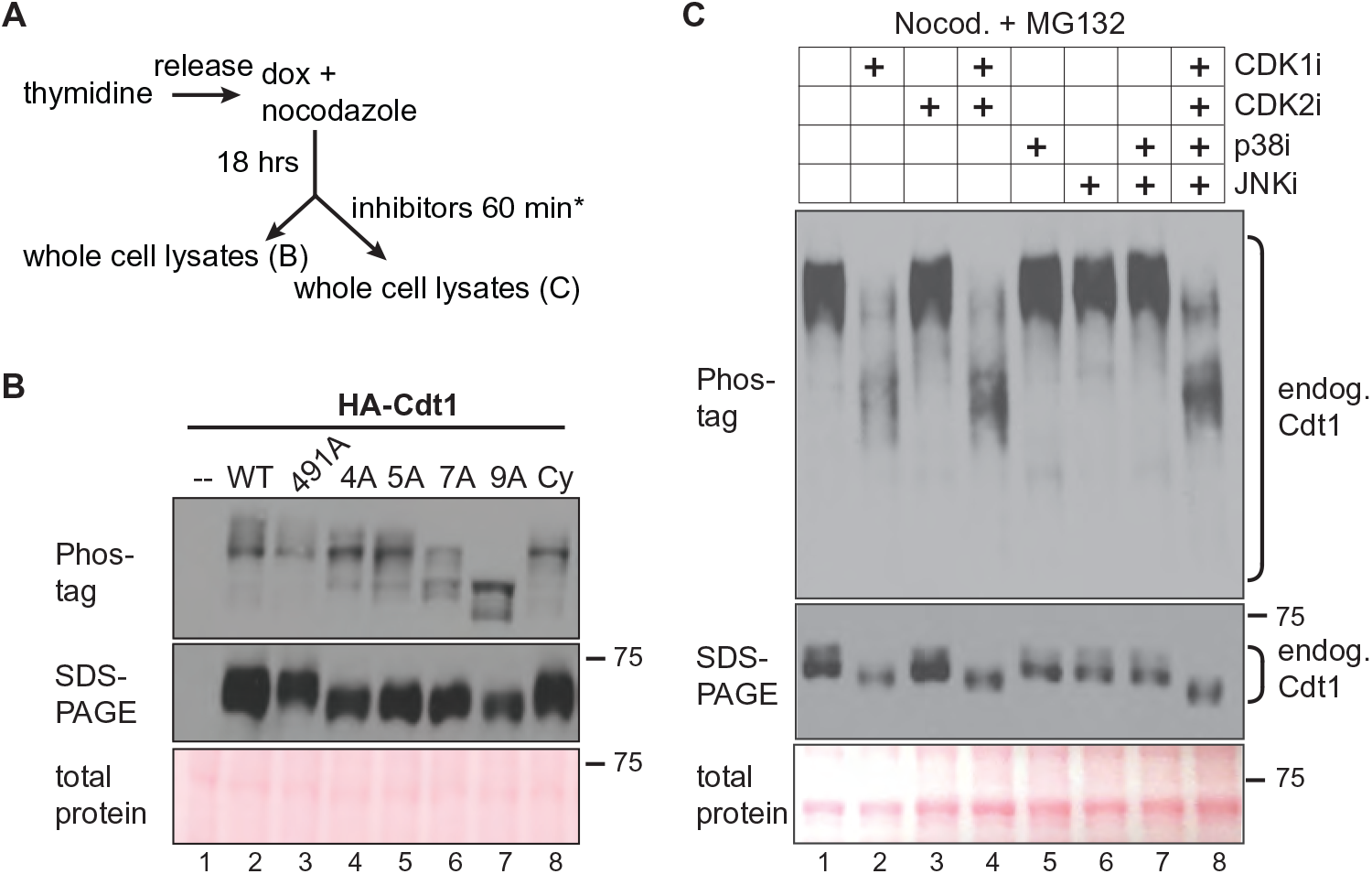
Cdt1 hyperphosphorylation is dependent on linker sites and CDK1 activity. **A)** Workflow for cell line synchronization and inhibitor treatment. **B)** Whole cell lysates were separated by Phos-tag SDS-PAGE (top) or standard SDS-PAGE (middle) followed by immunoblotting for ectopic Cdt1 (HA); total protein stain serves as a loading control. **C)** Cells were synchronized with nocodazole as in A, then mock treated or treated with 10 µM RO-3306 (lane 2), 6 µM CVT313 (lane 4), 30 µM SB203580 (lane 5), 10 µM JNK inhibitor VIII (lane 6), or combinations of inhibitors as indicated for 1 hour except that RO3306 treatment was for only the final 15 minutes to preserve mitotic cell morphology. All cells were simultaneously treated with 20 μM MG132 to prevent premature mitotic exit. Endogenous Cdt1 phosphorylation was assessed by standard or Phos-tag SDS-PAGE followed by immunoblotting; total protein stain serves as a loading control. The example shown is representative of more than three independent experiments.

Compared to Cdt1-5A, the Cdt1-7A variant had only slightly shifted distribution towards faster migration on Phos-tag gels (Fig. 3B, compare lanes 5 and 6). Cdt1-4A and the Cdt1-Cy mutant migrated on Phos-tag gels with a pattern very similar to Cdt1-5A whereas Cdt1-S491A migration was indistinguishable from Cdt1-WT (Fig. 3B, lanes 2-5 and 8). Cdt1-9A from synchronized cells migrated even faster than Cdt1-7A on Phos-tag gels, demonstrating that one or both T29 and S31 are also phosphorylated after S phase. Moreover, the difference in migration between Cdt1-9A and Cdt1-Cy suggests that either some residual kinase binding remains in the Cy motif mutant or that non-Cy-dependent kinases can phosphorylate some of the sites mutated in Cdt1-9A. These patterns indicate that Cdt1 is multiply phosphorylated late in the cell cycle on a collection of sites that includes both the N-terminal CDK sites and importantly, the set of linker phosphorylation sites that inhibit Cdt1 activity and restrict re-replication.

### Cyclin A/CDK1 is the primary Cdt1 kinase during G2 and M phases

To determine which kinase(s) is responsible for Cdt1 phosphorylation, we assessed the effects of kinase inhibitors. As a first step, we analyzed the migration of *endogenous* Cdt1 on Phos-tag gels using lysates from asynchronously proliferating or synchronized cells. Cdt1 from asynchronous cells migrates primarily as two bands on Phos-tag gels, and both forms are absent from lysates of UV-irradiated cells. Cdt1 is degraded during repair of UV-induced damage [45–48], so we conclude that both bands are endogenous Cdt1. Endogenous Cdt1 in nocodazole-synchronized cells migrated as a tight set of very slow-migrating species that are converted to the two faster forms by phosphatases in vitro prior to gel electrophoresis (Supplementary Fig S4A, lanes 3 and 4).

We then synchronized cells in nocodazole to induce maximal Cdt1 phosphorylation and tested the effects of pharmacological kinase inhibitors on the migration of endogenous Cdt1 using Phos-tag gels. All nine of the sites we had altered are predicted to be potential targets of both CDKs and MAPKs since all nine are serine or threonine followed by proline [49–52](Supplementary Fig. S1). Both kinase classes are active in G2 [53–55], so we postulated that during normal G2 and M phases these Cdt1 sites are phosphorylated by CDK and/or MAPK. In addition to the kinase inhibitors, we also co-treated with the proteasome inhibitor MG132 to prevent cyclin or other ubiquitin-mediated protein degradation. We first treated nocodazole-arrested cells with inhibitors of p38 or JNK, two stress-activated MAP kinases which we previously showed can phosphorylate the linker region during a stress response [17] (p38 inhibitor SB203580 and c-Jun N-terminal kinase JNK inhibitor VIII). These MAPK inhibitors, either alone or in combination, had no effect on mitotic Cdt1 migration on Phos-tag gels (Fig. 3C, lanes 5-7, compared to lane 1). We confirmed that the inhibitors were active in these cells at these concentrations by analyzing known downstream substrates (Supplementary Fig. S4B-D) [17, 56, 57]. We also tested inhibitors of CDK1 and CDK2 singly or in combination. In contrast to the effects of MAPK inhibitors, the slow migration of phospho-Cdt1 was largely reversed by treatment with CDK1 inhibitor RO-3306 [58] for just 15 minutes (Fig. 3C, compare lanes 2 and 4 to lane 1, treatment was shorter to preserve mitotic cell morphology), but not when treated with the CDK2 inhibitor CVT313 for an hour, (Fig. 3C, lane 3).

CDK1 is normally activated by either Cyclin A or Cyclin B, and we next sought to identify which cyclin is responsible for directing CDK1 to phosphorylate Cdt1. We therefore took advantage of the polyhistidine tag at the C-terminus of the Cdt1-WT construct to retrieve Cdt1 from lysates of transiently transfected, nocodazole-arrested cells. As a control, we included the Cdt1-Cy variant with a disrupted cyclin binding motif [40]. We analyzed Cdt1-bound proteins from these lysates for the presence of endogenous cyclin and CDK subunits. Cdt1-WT interacted with both CDK1 and CDK2, and strongly interacted with Cyclin A, but not at all with either Cyclin B or Cyclin E (Fig. 4A). Cdt1-Cy retrieved no cyclins or CDKs, indicating that the only strong CDK binding site in Cdt1 is the RRL at positions 66-68. Since Cdt1 binds Cyclin A, CDK1, and CDK2, but inhibiting CDK1 and not CDK2 affected Cdt1 phosphorylation in nocodazole-arrested, we conclude that Cyclin A/CDK1 is responsible for the inactivating Cdt1 phosphorylations during G2 and M phases. Cyclin A/CDK2 also binds Cdt1 and contributes to Cdt1 degradation during S phase [34, 39, 40], but our results indicate that in nocodazole-arrested cells, CDK2 activity is not required for Cdt1 phosphorylation.

**Figure 4.**
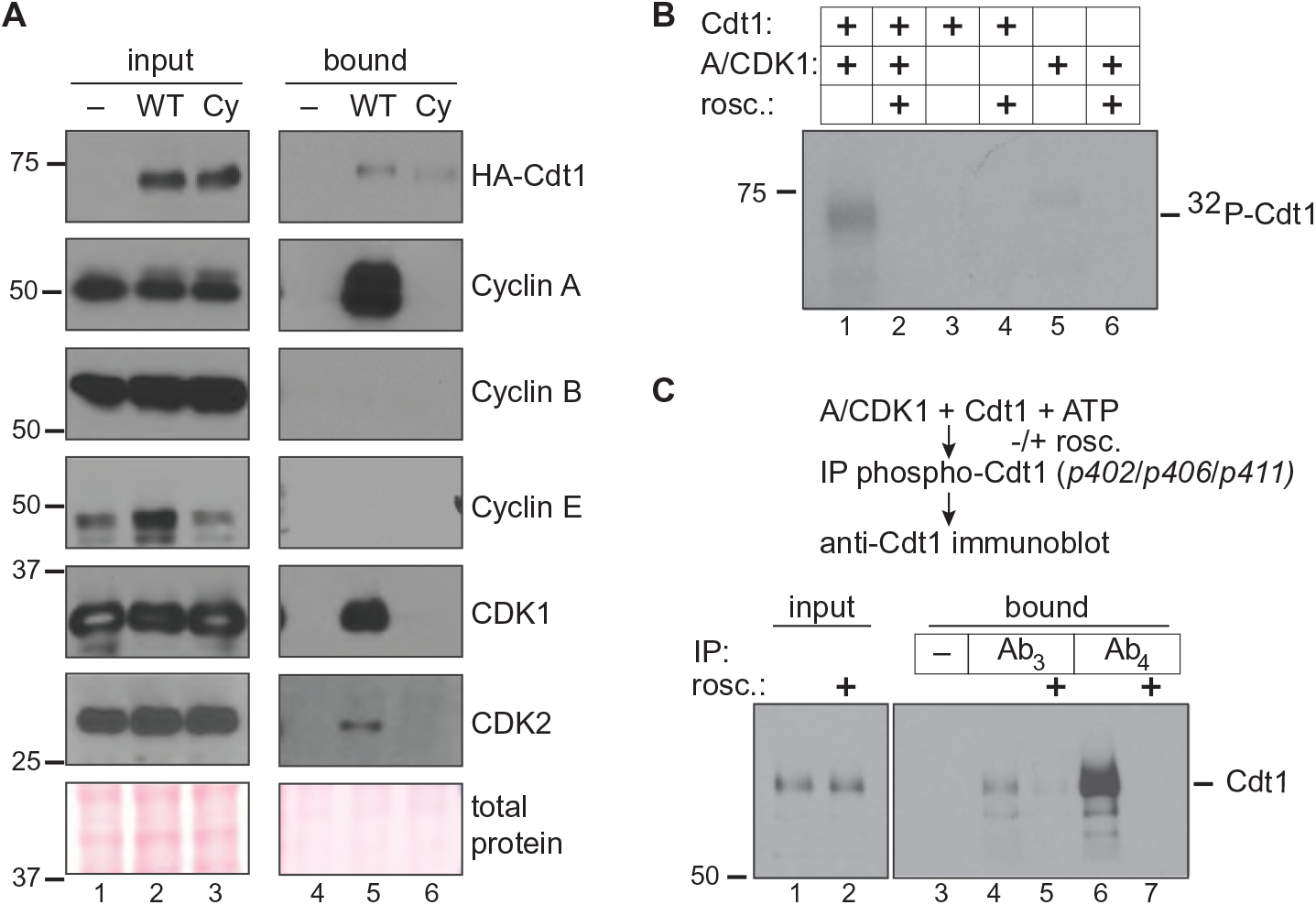
Cyclin A/CDK1 phosphorylates Cdt1 linker sites. **A)** HEK 293T cells were transfected with control plasmid or plasmid producing His-tagged Cdt1-WT or a Cdt1-variant that cannot bind CDKs (Cdt1-Cy) then synchronized with nocodazole and harvested by mitotic shake off. Cdt1 was retrieved on nickel-agarose, and the indicated endogenous proteins were detected in whole cell lysates (lanes 1-3) and bound fractions (lanes 4-6) by immunoblotting. The result is representative of at least two independent experiments. **B)** Recombinant partially-purified Cdt1 was incubated with purified Cyclin A/CDK1 in the presence of ^32^P-γ-ATP in kinase buffer for one hour at 30°C. Control reactions contained Cdt1 only, kinase only, or were complete reactions in the presence of 20 µM roscovitine (CDK inhibitor) as indicated. Reactions were separated by SDS-PAGE followed by autoradiography. **C)** Recombinant Cdt1 was incubated with purified Cyclin A/CDK1 in the presence of unlabeled ATP as in B; roscovitine was included as indicated. Reactions were subjected to immunoprecipitation with either pre-immune serum or immune sera to retrieve Cdt1 phosphorylated at S402, S406, and T411; Ab_3_ and Ab_4_ are consecutive test bleeds from the immunized rabbit. Both input and bound proteins were probed for total Cdt1 by immunoblotting.

To determine if Cyclin A/CDK1 can directly phosphorylate Cdt1, we incubated Cdt1 that had been partially purified from transfected cells with purified Cyclin A/CDK1 and [γ-^32^P]-ATP. Cdt1 was directly phosphorylated *in vitro* by Cyclin A/CDK1, and this phosphorylation was blocked by the general CDK1/CDK2 inhibitor, roscovitine (Fig. 4B, lanes 1 and 2). However, this assay does not distinguish between phosphorylation at the previously studied N-terminal CDK target sites, T29 and S31, and sites in the linker region or elsewhere. To test specifically for linker region phosphorylations, we repeated the *in vitro* kinase reactions in the presence of unlabeled ATP and then subjected the reactions to immunoprecipitation with a phospho-specific antibody raised against Cdt1 sites S402, S406 and T411. We had previously described this antibody as being suitable for immunoprecipitation (though not for immunoblotting) [17]. Two different test sera from that antibody production detect Cdt1 phosphorylation by immunoprecipitation followed by immunoblotting with a general Cdt1 antibody; these sera are labelled Ab_3_ and Ab_4_. By this method, we detected direct Cyclin A/CDK1-mediated Cdt1 phosphorylation at the inhibitory linker sites *in vitro* (Fig. 4C, lanes 4 and 6).

### Cdt1 phosphorylation blocks MCM binding

We next explored the molecular mechanism of Cyclin A/CDK1-mediated Cdt1 licensing inhibition. The inhibitory phosphorylation sites are not visible in any currently available Cdt1 atomic structures. Nonetheless, our homology model of the human Cdt1-MCM complex ([28] and Fig. 5A) led us to speculate that phosphorylation-induced changes at this linker could affect MCM binding. We first compared the MCM binding ability of Cdt1-WT to the Cdt1-Cy variant that cannot bind Cyclin A/CDK1. We transiently transfected cells with these plasmids and then immunoprecipitated MCM2 from asynchronously growing cells or from cells arrested in nocodazole. MCM6 serves as a marker of the MCM complex retrieved by the MCM2 immunoprecipitation. Asynchronously growing cells spend more time in G1 than in G2 and have mostly hypophosphorylated Cdt1. Thus as expected, there was little difference in MCM binding ability between Cdt1-WT and Cdt1-Cy in asynchronous cells (Fig. 5B, lanes 6 and 7). In contrast, in nocodazole-arrested cells where Cdt1-WT was hyperphosphorylated, but Cdt1-Cy was less phosphorylated, the Cdt1-Cy variant bound MCM significantly better than Cdt1-WT (Fig. 5B, lanes 9 and 10). This difference in binding was independent of the presence of high Geminin levels in mitotic cells (Fig. 5B, lanes 3 and 4) which is also known to affect Cdt1-MCM binding [22, 23].

**Figure 5.**
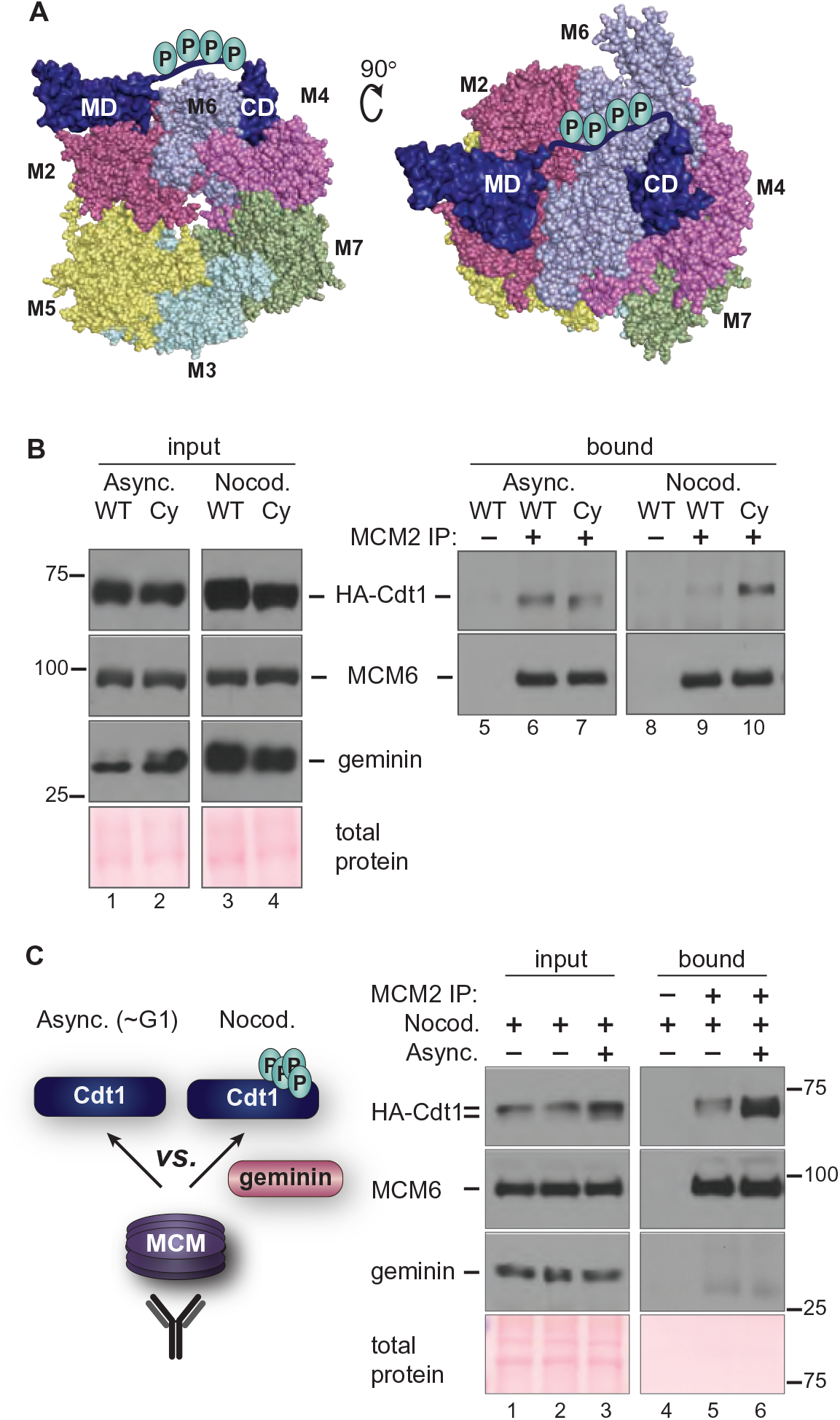
Hyperphosphorylation impairs Cdt1-MCM binding. **A)** Two views of a homology model of the human MCM_2-7_-Cdt1 complex as described in Pozo *et al.* 2018; numbers refer to individual MCM subunits. The disordered linker containing phosphorylation sites is hand-drawn connecting the two structured Cdt1 domains (MD and CD) in the model. **B)** Asynchronously growing or nocodazole-arrested HEK293T cells ectopically expressing HA-tagged Cdt1-WT or the Cdt1-Cy variant were lysed and subjected to immunoprecipitation with anti-MCM2 antibody. Whole cell lysates (lanes 1-4) and bound proteins (lanes 5-10) were probed for HA, MCM6 and Geminin, respectively; total protein stain serves as a loading control. The results are representative of two independent experiments. **C)** A lysate of nocodazole-arrested (Cdt1 hyperphosphorylated, Geminin-expressing) U2OS cells producing HA-tagged Cdt1-WT was mixed with lysate from the same cells growing asynchronously as indicated. Asynchronous cells contain mostly hypophosphorylated Cdt1 and very little Geminin. These lysates were then subjected to immunoprecipitation with anti-MCM2 antibody and probed for bound Cdt1. Input lysates (lanes 1-3) and bound proteins (lanes 4-6) were probed for HA-Cdt1, MCM6 (as a marker of the MCM complex), and Geminin. The example shown is representative of three independent experiments.

We then set out to test if MCM interacts with hyperphosphorylated G2 Cdt1 less well than with hypophosphorylated G1 Cdt1 (i.e. if phosphorylation impairs Cdt1-MCM binding). We noted however that simply comparing co-immunoprecipitations from lysates of G1 and G2 phase cells is complicated by the presence of the Cdt1 inhibitor, Geminin, which interferes with the Cdt1-MCM interaction and is only present in S and G2 cells. Because Geminin is differentially expressed in G1 and G2 cells, the comparison would not be fair. To account for the effects of Geminin, we prepared a lysate of asynchronously-proliferating, mostly G1 cells, then mixed this lysate with lysate from nocodazole-arrested cells that contains both Geminin and hyperphosphorylated Cdt1 (Fig. 5C, lane 3 input). In this way, we created a similar opportunity for MCM to bind either hyper- or hypophosphorylated Cdt1. We then immunoprecipitated endogenous MCM2 and probed for MCM6 as a marker of the MCM complex and for Cdt1. For comparison, we immunoprecipitated MCM2 from an unmixed lysate of nocodazole-arrested cells with only hyperphosphorylated Cdt1. As expected, Geminin did not co-precipitate with MCM since the Cdt1-Geminin and Cdt1-MCM interactions are mutually exclusive (Fig. 5C). Importantly, we found that Cdt1 bound by the MCM complex in the mixed lysates was enriched for the faster migrating hypophosphorylated Cdt1 relative to hyperphosphorylated Cdt1. Moreover, the total amount of Cdt1 bound to MCM was much higher when hypophosphorylated Cdt1 was available than when the only form of Cdt1 was hyperphosphorylated (Fig. 5C, compare lanes 5 and 6). This preferential binding suggests that Cdt1 phosphorylation disrupts interaction with the MCM complex, and that this disruption contributes to re-replication inhibition in G2 and M phases. We note that this is the first example of direct regulation of the Cdt1-MCM interaction by post-translational modification.

### Cdt1 dephosphorylation at the M-G1 transition requires PP1 phosphatase activity

Our finding that Cdt1 phosphorylation in G2 and M phase inhibits its ability to bind MCM suggests that Cdt1 must be dephosphorylated in the subsequent G1 phase to restore its normal function. To explore this notion, we first monitored Cdt1 expression and phosphorylation in cells progressing from M phase into G1. We released nocodazole-arrested cells and collected time points for analysis by immunoblotting (Fig. 6A). Like the mitotic cyclins, Geminin is a substrate of the Anaphase Promoting Complex/Cyclosome (APC/C) [59], and as expected for an APC/C substrate, Geminin was degraded within 60 minutes of mitotic release. In contrast, Cdt1 was not degraded during the M-G1 transition but rather, was rapidly dephosphorylated coincident with Geminin degradation (Fig. 6A, compare lanes 3 and 4). We next investigated which phosphatase is required for Cdt1 dephosphorylation. We first tested phosphatase inhibitors for the ability to prevent Cdt1 dephosphorylation after CDK1 inhibition. We tested inhibitors of protein phosphatase 1 (PP1) and protein phosphatase 2A (PP2A), and these two families account for the majority of protein dephosphorylation in cells [60]. We treated nocodazole-arrested cells with the CDK1 inhibitor to induce Cdt1 dephosphorylation in the presence or absence of calyculin A (Cal A) or okadaic acid (OA) [61]. Both compounds are potent inhibitors of both PP1 and PP2A, but calyculin A is more effective than okadaic acid for inhibiting PP1, particularly at the concentrations we tested [62]. We found that calyculin A preserved Cdt1 hyperphosphorylation (Fig. 6B, compare lanes 2 and 3) whereas low concentrations of okadaic acid that inhibit PP2A but not PP1 did not affect Cdt1 dephosphorylation (Supplementary Fig. S5). In addition, we released nocodazole-arrested cells into G1 phase for 30 minutes (to initiate mitotic progression) and then treated the cells with calyculin A. As a control, we probed for MCM4, a known PP1 substrate that is normally dephosphorylated in G1 phase [63]; calyculin A prevented MCM4 dephosphorylation (Fig. 6C). PP1 inhibition also largely prevented Cdt1 dephosphorylation during the mitosis-G1 phase transition without blocking overall mitotic progression as evidenced by Geminin degradation (Fig. 6C, lanes 2 and 3). These results suggest that a PP1 family phosphatase is required for Cdt1 dephosphorylation. By extension, we suggest that PP1 activity is required to re-activate Cdt1-MCM binding and origin licensing in G1 phase.

**Figure 6.**
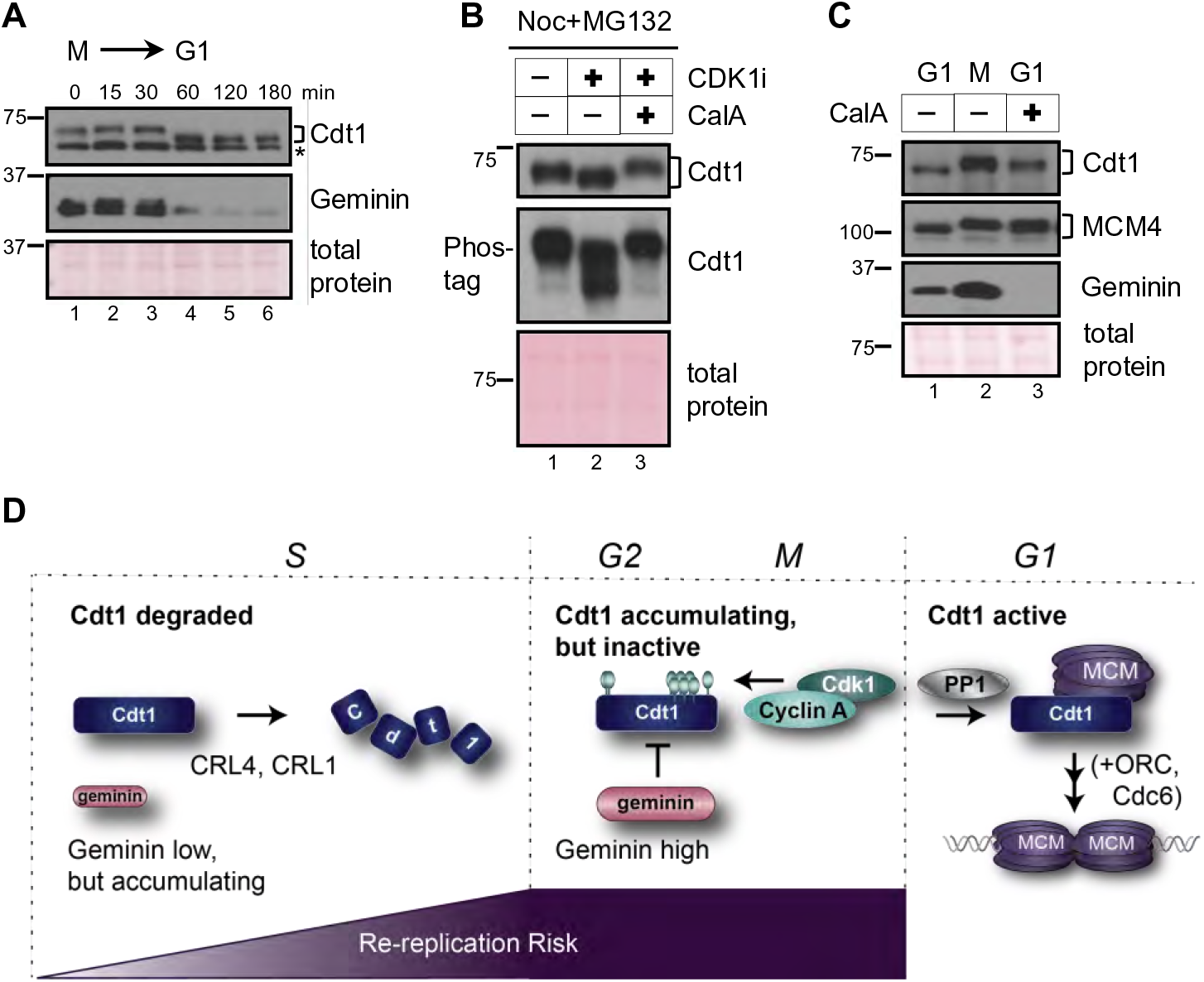
Cdt1 dephosphorylation at the M-G1 transition requires PP1. **A)** Nocodazole-arrested U2OS cells were released into fresh medium and collected at the indicated time points. Endogenous Cdt1 phosphorylation (top) and Geminin (middle) degradation were analyzed by immunoblotting; Ponceau S staining for total protein and a non-specific band (*) serve as loading controls. The results are representative of two independent experiments. **B)** Nocodazole-arrested U2OS cells were mock treated (lane 1) or treated with 10 µM RO-3306 (CDK1i, lane 2), or treated with both 10 µM RO-3306 and with 20 nM calyculin A as indicated (CalA, lane 3). Endogenous Cdt1 phosphorylation was analyzed by standard or Phos-tag SDS-PAGE followed by immunoblotting; total protein stain serves as a loading control. The results are representative of three independent experiments. **C)** Nocodazole-arrested U2OS cells (lane 2) were released into fresh medium for 3 hours and mock treated (lane 1) or treated with 20 nM calyculin A 30 minutes after release (lane 3). Endogenous Cdt1 or MCM4 phosphorylation and total Geminin were detected by immunoblotting; total protein stain serves as a loading control. The results are representative of three independent experiments. **D)** Model. In S phase Cdt1 is targeted for degradation, first by the CRL4^Cdt2^ E3 ubiquitin ligase at the onset of S phase and then additionally by SCF^Skp2^ after phosphorylation by Cyclin A/CDK2. Geminin accumulates starting in early S phase. The amount of duplicated DNA at risk of re-replication is lowest in early S and highest in G2. In late S and G2 phase Cdt1 re-accumulates and Geminin is at high levels. Cyclin A/CDK1 phosphorylates Cdt1, and both Geminin and Cdt1 hyperphosphorylation independently block Cdt1-MCM binding. At the M→G1 transition Protein Phosphatase 1 is required for Cdt1 dephosphorylation to reactivate MCM loading by Cdt1, ORC, and Cdc6.

## Discussion

### Cell cycle-dependent Cdt1 phosphorylation

Metazoan Cdt1 is degraded during S phase, and this degradation is essential to prevent re-replication [34, 40, 64, 65]. Perhaps counter-intuitively, Cdt1 then actively accumulates beginning in late S phase, and by mitosis reaches a level similar to Cdt1 in G1 phase [16–21]. Despite the potential risk for re-licensing and re-replicating G2 DNA, these high Cdt1 levels serve two purposes: 1) Cdt1 is essential for stable kinetochore-microtubule attachments [21, 66], and 2) high levels of Cdt1 in mitosis can improve licensing efficiency in the next G1 phase [18]. In this study, we discovered that Cdt1 phosphorylation during G2 phase inhibits Cdt1 licensing activity and contributes to preventing DNA re-replication during the time that Cdt1 levels are high in G2 and M phase.

We analyzed a cluster of Cyclin A/CDK1-dependent phosphorylation sites that are distinct from the previously characterized CDK sites at T29 and S31. This region of Cdt1 is not strongly conserved among vertebrates (Fig. 1A and Supplementary Fig. S1), but most vertebrate Cdt1 linker sequences are nonetheless predicted to be similarly disordered, and most have at least one candidate CDK phosphorylation site (Supplementary Fig. S1). Interestingly, altering two additional sites in this region did not exacerbate the re-replication phenotype suggesting that four phosphorylations are sufficient to achieve maximal human Cdt1 inhibition. In that regard, multisite Cdt1 linker phosphorylation may resemble other examples of cell cycle-dependent multisite phosphorylation in which the total negative charge is more important than the specific phosphorylated position [67].

We show that Cdt1 only binds strongly to endogenous Cyclin A and binds neither Cyclin E nor Cyclin B. The fact that Cdt1 is unlikely to be a direct target of Cyclin E activity is reassuring since Cyclin E is active in late G1 phase at the same time that MCM is loading, and it would be counterproductive to inhibit Cdt1 activity in late G1. On the other hand, undetectable Cyclin B binding is somewhat surprising since Cdt1 remains phosphorylated throughout all of mitosis, and Cyclin A can be degraded at the beginning of mitosis [68, 69]. It may be that Cdt1 phosphorylation is maintained throughout mitosis by the high levels of active Cyclin B/CDK1 without the need for tight CDK-Cdt1 binding, or that a residual amount of tightly-bound Cyclin A maintains Cdt1 phosphorylation, or that some unknown cellular kinase or phosphatase inhibitor keeps Cdt1 phosphorylated even after Cyclin A is degraded. If a minor kinase takes over from Cyclin A/CDK1, its activity is clearly also lost after treatment with a relatively selective CDK1 inhibitor. We also acknowledge that in actively proliferating cells Cyclin A/CDK2 could contribute to direct Cdt1 inactivation in late S and G2 phase in a time window after Cdt1 accumulation but before substantial Cyclin A/CDK1 activation, but our synchronization experiment did not detect a role for CDK2 activity.

We demonstrate here that the CDK docking motif at Cdt1 positions 68-70 is required for phosphorylation not only at the previously investigated T29 position, but also at sites more than 300 residues towards the Cdt1 C-terminus. The structure of the yeast Cdt1-MCM complex indicates that when bound to MCM, Cdt1 is in a relatively extended conformation with the linker quite distant from the N-terminal domain [70–72]. We did not model the N-terminal domain of human Cdt1 because it bears almost no sequence similarity to the corresponding domain of budding yeast Cdt1. Our discovery that the mammalian Cy motif controls phosphorylation at sites very distant in the primary sequence prompt speculation that Cdt1 in isolation from MCM may adopt a conformation with the linker relatively close to the N-terminal regulatory domain for phosphorylation by the Cy motif-bound Cyclin A/CDK1.

### Phosphorylation inhibits Cdt1 binding to MCM

We found that hyperphosphorylated Cdt1 binds MCM poorly relative to hypophosphorylated Cdt1. This observation provides a simple mechanism for Cyclin A/CDK1-mediated phosphorylation to inhibit Cdt1 licensing activity. Both the Cdt1 N-terminal domain and the linker region are predicted to be intrinsically disordered, and the fact that these regions were excluded from mammalian Cdt1 fragments subjected to structure determination supports that prediction [26, 27]. The only structure of full-length Cdt1 available to date is a component of the budding yeast Cdt1-MCM or ORC/Cdc6/Cdt1/MCM complexes [71, 72], and budding yeast Cdt1 lacks candidate phosphorylation sites in the linker region. For this reason, we cannot determine precisely how phosphorylation in the linker inhibits MCM binding. We suggest however, that the introduction of multiple phosphorylations either induces a large conformational change in Cdt1 that prevents it from extending around the side of the MCM ring or alternatively, these phosphorylations may repel Cdt1 from the MCM surface (Fig. 5A).

We had previously established that the p38 and JNK stress-activated MAP kinases can phosphorylate at least some of these same inhibitory sites in Cdt1 [17], and a separate study reported a subset of these plus additional sites as potential JNK targets [41]. Both p38 and JNK are active during a G2 arrest [53–55, 73, 74], but our inhibitor results indicate that Cyclin A/CDK1 is dominant for Cdt1 phosphorylation during G2 and M phases in these cells. On the other hand, our findings here also shed light on the molecular mechanism of stress-induced origin licensing inhibition [17]. We postulate that stress MAPK-mediated Cdt1 hyperphosphorylation at the linker region blocks Cdt1-MCM binding in stressed G1 cells to prevent origin licensing. This phosphorylation blocks initial origin licensing by the same mechanism that prevents origin re-licensing in G2 and M phases. The p38 MAPK family is also active in quiescent cells [53, 54], and Cdt1 from lysates of serum-starved cells has slower gel mobility reminiscent of the same shift we and other observe in G2 and M phase cells [17]. We thus speculate that Cdt1 in quiescent cells is inhibited by a similar mechanism as the one we defined here.

The nine phosphorylation sites we tested in this study are certainly not the only phosphorylation sites in human Cdt1. Unbiased phosphoproteomics studies have detected phosphorylation at a total of 22 sites, 13 of which are also S/T-P sites [38]. In addition, a domain in the N-terminal region restrains Cdt1 licensing activity by influencing chromatin association includes at least two other mitotic CDK/MAPK sites [19]. It is not known if phosphorylation at those sites is strictly cell cycle-dependent or requires the Cy motif. The fact that Cdt1-Cy has the highest activity of all the variants tested here may be a reflection of that additional negative regulation in the so-called “PEST domain.” Alternatively, the Cy motif mutation may disrupt more than only Cyclin binding such as has been recently reported for ORC [75]. Clearly the spectrum of Cdt1 biological activities can be tuned by combinations of phosphorylations and dephosphorylations, and continued in-depth analyses will yield additional insight into Cdt1 regulation and function.

Approximately one-third of all eukaryotic proteins may be dephosphorylated by PP1 [60]. PP1 binds some of its substrates directly via a short motif, RVxF, KGILK or RKLHY [60, 76]. Human Cdt1 contains several such candidate PP1 binding motifs and thus may be a direct target of PP1. Alternatively, Cdt1 dephosphorylation may require an adapter to bind PP1 similar to the role of the Rif1 adapter for MCM dephosphorylation [63, 77, 78]. In either case, the fact that hyperphosphorylated Cdt1 binds MCM poorly, plus the fact that the levels of Cdt1 do not change from M phase to G1 (i.e. Cdt1 is not degraded and resynthesized at the M-G1 transition), means that PP1-dependent Cdt1 dephosphorylation *activates* origin licensing. In that regard, dephosphorylation is the first example of direct Cdt1 activation, and it complements the indirect activation by Geminin degradation at the M to G1 transition.

### A sequential relay of re-replication inhibition mechanisms

We propose that Cdt1 activity is restricted to only G1 through multiple regulatory mechanisms during a single cell cycle, but that the relative importance of individual mechanisms changes at different times after G1 (Fig. 6D). At the onset of S phase Cdt1 is first subjected to rapid replication-coupled destruction via CRL4^Cdt2^ which targets Cdt1 bound to DNA-loaded PCNA [79]. This degradation alone is not sufficient to prevent re-replication however, and a contribution from Cyclin A/CDK2 to create a binding site for the SCF^Skp2^ E3 ubiquitin ligase is also essential [34]. We suggest that SCF^Skp2^-targeting occurs primarily in mid and late S phase based on the dynamics of Cyclin A accumulation. A reinforcing mechanism for Cdt1 degradation is more important in mid and late S phase than in early S phase because the amount of DNA that has already been copied increases throughout S phase. The consequences of licensing DNA that hasn’t yet been copied are presumably benign, but as S phase proceeds, the amount of DNA that has been copied already (i.e. the substrate for re-replication) also increases. The Cdt1 inhibitor, Geminin, begins to accumulate near the G1-S transition, and its levels increase along with the amount of replicated DNA until Geminin is targeted for degradation by the APC/C during mitosis [24, 59]. Geminin binding to Cdt1 interferes with Cdt1-MCM binding, and since Cdt1-MCM binding is essential for MCM loading, Geminin prevents re-licensing [35, 36]. This inhibition is particularly important once Cdt1 re-accumulates after S phase is complete [37]. Just as CRL4^Cdt2^-mediated degradation in S phase is not sufficient to fully prevent re-replication, we demonstrated that the presence of Geminin alone is not sufficient to inhibit Cdt1 during G2. Cdt1 phosphorylation in a linker domain between two MCM binding sites also prevents Cdt1-MCM binding. These (and potentially more) mechanisms to restrain Cdt1 activity are also reinforced by regulation to inhibit ORC, Cdc6, PR-Set7, and other licensing activators [4, 33, 80, 81]. The relative importance of any one mechanism will be influenced by cell type and species. Given that there are many thousands of origins in mammalian genomes, and the consequences of even a small amount of re-replication are potentially dire, we suggest that precise once-and-only-once replication requires that Cdt1 be inhibited by at least two mechanisms at all times from G1 through mitosis.

## Materials and Methods

### Cell Culture and Manipulations

U2OS Flp-in Trex cells [82] bearing a single FRT site (gift of J. Aster) and HEK 293T cells were arrested by thymidine-nocodazole synchronization by treatment with 2 mM thymidine for 18 h followed by release into 100 nM nocodazole for 10 h. Cells were treated with inhibitors for 1 hour and harvested by mitotic shake-off, with the exception that RO-3306 treatment was for just 15 minutes. Cells were treated with 10 μM, RO-3306 (Sigma), 6 μM CVT313 (Sigma), 10 μM JNK inhibitor VIII (Sigma), 30 μM SB203580 (Sigma), 20 μM MG132 (Sigma), Okadaic acid (Abcam #ab120375), or 20 nM, calyculin A (LC Laboratories) as indicated. HEK 293T cells were transfected with Cdt1 expression plasmids using PEI Max (Sigma) and cultured for 16 hours. All cell lines were validated by STR profiling and monitored by mycoplasma testing. For flow cytometry, cells were cultured in complete medium with 1 μg/mL doxycycline for 48 hours labeled with 10 μM EdU (Sigma) for 1 hour prior to harvesting.

### Antibodies

Antibodies were purchased from the following sources: Cdt1 (Cat# 8064), Chk1 (Cat# 2345), phospho-Chk1 S345 (Cat# 2341), Cyclin E1 (Cat#4129), MAPKAPK-2 (Cat#), Phospho-MAPKAPK-2 T334 (Cat#3007), phospho-Histone H2A.X Ser139 (Cat#9718) from Cell Signaling Technologies; hemagglutinin (HA) (Cat#11867423001) from Roche; Geminin (Cat#sc-13015), Cdc6 (Cat#sc-9964), MCM6 (Cat#sc-9843), Cyclin A (Cat#sc-596), Cyclin B1 (Cat#sc-245) and CDK2 (Cat#sc-163) from Santa Cruz Biotechnology; MCM4 (Cat#3728) from Abcam. MCM2 antibody (Cat#A300-191A) used for co-immunoprecipitation experiment was purchased from Bethyl Laboratories. MCM2 antibody (BD Biosciences, Cat#610700) was used for analytical flow cytometry. Serum to detect CDK1 was a gift from Y. Xiong (University of North Carolina), and MPM2 antibody was a gift from R. Duronio [83] (University of North Carolina). The phosphospecific Cdt1 antibody was described in Chandrasekaran et al [17].; the third and fourth test bleeds are active for Cdt1 immunoprecipitation. Alexa 647-azide and Alexa-488-azide used in flow cytometry analyses was purchased from Life Technologies, and secondary antibodies for immunoblotting and immunofluorescence were purchased from Jackson ImmunoResearch.

### Protein-protein interaction assays

For polyhistidine pulldown assays, cells were lysed in lysis buffer (50 mM HEPES pH 8.0, 33 mM KAc, 117 mM NaCl, 20 mM imidazole, 0.5% triton X-100, 10% glycerol) plus protease inhibitors (0.1 mM AEBSF, 10 µg/mL pepstatin A, 10 µg/mL leupeptin, 10 µg/mL aprotinin), phosphatase inhibitors (5 µg/mL phosvitin, 1 mM β-glycerol phosphate, 1 mM Na-orthovanadate), 1 mM ATP, 1 mM MgC_l2_, 5 mM CaCl_2_ and 15 units of S7 micrococcal nuclease (Roche). Lysates were sonicated for 10 seconds at low power followed by incubation on ice for 30 minutes and clarification by centrifugation at 13,000 x g for 15 minutes at 4°C. The supernatants were incubated with nickel NTA agarose beads (Qiagen) for 2 hours at 4°C with rotation. Beads were rinsed 4 times rapidly with ice-cold lysis buffer followed by boiling in SDS sample buffer for 5 minutes prior to immunoblot.

For co-immunoprecipitation assays, cells were lysed in Co-IP buffer (50 mM HEPES pH 7.2, 33 mM KAc, 1 mM MgCl_2_, 0.5% triton X-100, and 10% glycerol) containing protease inhibitors (0.1 mM AEBSF, 10 µg/mL pepstatin A, 10 µg/mL leupeptin, 10 µg/mL aprotinin), phosphatase inhibitors (5 µg/mL phosvitin, 1 mM β-glycerol phosphate, 1 mM Na-orthovanadate), 1 mM ATP, and supplemented with 5 mM CaCl2 and 15 units of S7 micrococcal nuclease (Roche). Lysates were sonicated for 10 seconds at low power followed by incubation on ice for 30 minutes and clarification by centrifugation at 13,000 x g for 15 minutes at 4°C. The supernatants were incubated and rotated with Protein A beads (Roche) with an anti-Mcm2 antibody (Bethyl, 1:1000) at 4°C with rotation for 4 hours. Beads were rinsed three times with ice-cold co-IP buffer then eluted by boiling in sample buffer for subsequent immunoblot analysis.

### Immunofluorescence microscopy

U2OS cells cultured on cover glass were fixed with 4% PFA for 15 minutes and permeabilized with 0.5% Triton in PBS for 5 minutes. Cells were blocked in 1% BSA for 30 minutes followed by incubation with primary antibody overnight at 4°C and secondary antibody for 1 hour at room temperature. Cells were stained with 1 μg/ml DAPI for 5 minutes before mounting with the ProLong® Gold Antifade mounting medium (life technologies). Fluorescent images were captured on a Nikon 2000E microscope. The areas of nuclei were measured by using the Adobe Photoshop software.

### Analytical flow cytometry

For cell cycle analysis, cells were cultured in complete medium with 1 ug/ml doxycycline for 48 hours. Cells were pulse labeled with 10 µM EdU (Sigma) for 60 minutes prior to harvesting by trypsinization. Cells were washed with PBS and then fixed in 4% paraformaldehyde (Sigma) followed by processing for EdU conjugation to Alexa Fluor 647-azide (Life Technologies). Samples were centrifuged and incubated in PBS with 1 mM CuSO_4_, 1 mM fluorophore-azide, and 100 mM ascorbic acid (fresh) for 30 min at room temperature in the dark then washed with PBS. Total DNA was detected by incubation in 1 µg/mL DAPI (Life Technologies) and 100 µg/mL RNAse A (Sigma).

For MCM loading analysis, U2OS cells were cultured in complete medium with 0.05 μg/mL doxycycline for 24 hours to induce expression of ectopic constructs.. Approximately 20% of this suspension was reserved for subsequent immunoblotting analysis while the remaining 80% was analyzed for bound MCM as described in Matson et al. [42]. Briefly, cells were extracted in cold CSK buffer (10 mM Pipes pH 7.0, 300 mM sucrose, 100 mM NaCl, 3 mM MgCl_2_) supplemented with 0.5% triton X-100, protease inhibitors (0.1 mM AEBSF, 1 µg/mL pepstatin A, 1 µg/mL leupeptin, 1 µg/mL aprotinin), and phosphatase inhibitors (10 µg/mL phosvitin, 1 mM β-glycerol phosphate, 1 mM Na-orthovanadate). Cells were washed with PBS plus 1% BSA and then fixed in 4% paraformaldehyde (Sigma) followed by processing for EdU conjugation. Bound MCM was detected by incubation with anti-MCM2 primary antibody at 1:200 dilution and anti-mouse-488 at 1:1,000 dilution at 37 °C for 1 hour. Data were collected on an Attune NxT flow cytometer (Thermo Fisher Scientific) and analyzed using FCS Express 7 (De Novo Software) software. Control samples were prepared omitting primary antibody or EdU detection to define thresholds of detection as in Matson et al 2017 [42].

### In vitro kinase assay

200 ng of recombinant human Cdt1 (OriGene, Cat #: TP301657) and 20 ng of purified Cyclin A/Cdk1 (Sigma cat. #CO244, lot SLBW3287) were incubated in kinase buffer (50 mM Tris pH 7.5, 10 mM MgCl_2_) supplemented with protease inhibitors (0.1 mM AEBSF, 10 µg/mL pepstatin A, 10 µg/mL leupeptin, 10 µg/mL aprotinin), phosphatase inhibitors (5 µg/mL phosvitin, 1 mM β-glycerol phosphate, 1 mM Na-orthovanadate), 10 µM ATP, 2 µCi of [γ-^32^P]-ATP, and in the presence or absence of roscovitine (20 µM) for 1 hr at 30°C. Reactions were stopped by adding loading buffer for subsequent SDS-PAGE and autoradiography.

### Statistical analysis

The differences were considered significant with a p-value less than 0.05. Values for multiple independent experiments were analyzed by one-way ANOVA for multiple comparisons without corrections (Fishers LSD test) but with pre-planned comparisons as described in the text. (Parallel analysis with Tukey’s multiple comparisons test did not alter interpretations.) Significance testing was performed using Prism 8 (GraphPad).

### Data availability

All expression constructs and data in the manuscript are available from the authors upon reasonable request.

## Supporting information

supplemental figures for Zhou, Pozo, et al.

## Acknowledgements

We thank Y. Xiong, R. Duronio, L. Graves, and J. Aster for the generous gifts of antibodies and reagents, and all members of the Cook lab for discussion and comments on the manuscript; S. Hailemariam and D. Tesfu contributed to early development steps in this project. We thank Jeffrey Jones for research support assistance. The UNC Hooker Imaging Core and the UNC Flow Cytometry Core Facility are supported in part by a National Institutes of Health Cancer Core Support Grant to the UNC Lineberger Comprehensive Cancer Center (CA016086). This work was also supported by National Institutes of Health grants F31GM121073 to P.N.P. and R01GM102413 and R01GM083024 to J.G.C.

## Author Contributions

Y.Z., S.O. and J.G.C. conceived the study. Y.Z. and P.N. P. performed and designed most of the experiments; H.M.S. performed the Cdt1 dephosphorylation experiments, S.O. generated some of the Cdt1 expression constructs and collected data in HeLa cells. J.G.C., Y.Z., and P.N.P. interpreted the results, analyzed the data, produced the figures, and wrote the manuscript.

---

**Figure S1. Cdt1 linker phosphorylation sites in 27 vertebrate sequences.**

A selection of 27 vertebrate sequences for comparison was taken from Miller *et al.* (2007), and Cdt1 protein sequences were retrieved from https://www.uniprot.org/. For the Cdt1 alignment, *Xenopus tropicalis* in Miller *et al.* was replaced with *Xenopus laevis* Cdt1, *Tupaia belangeri* was replaced with *Tupaia chinensis*, and no Cdt1 sequence for *Echinops telfairi* (tenrec) was available. These 27 full-length sequences were aligned with ClustalW at https://www.genome.jp/tools-bin/clustalw using the default settings, and the resulting alignment was visualized with BoxShade, 50% identity or similarity were shaded medium and light grey (https://embnet.vital-it.ch/software/BOX_form.html). The portion corresponding to the Cdt1 linker domain is shown using common names. All potential CDK/MAPK phosphorylation sites are shaded green, and an 85 residue insertion in chicken Cdt1 lacking any potential CDK/MAPK phosphorylation sites was deleted for clarity. The 27 sequences are from the following species: *Homo sapiens, Pan troglodytes, Macaca mulatta, Otolemur garnettii, Tupaia chinensis, Rattus norvegicus, Mus musculus, Cavia porcellus, Oryctolagus cuniculus, Sorex araneus, Erinaceus europaeus, Canis familiaris, Felis catus, Equus caballus, Bos Taurus, Dasypus novemcinctus, Loxodonta Africana, Monodelphis domestica, Ornithorhynchus anatinus, Gallus gallus, Anolis carolinensis, Xenopus laevis, Tetraodon nigroviridis, Takifugu rubripes, Gasterosteus aculeatus, Oryzias latipes,* and *Danio rerio*.

**Figure S2. Unphosphorylatable Cdt1 induces giant nuclei formation and DNA damage.**

**A)** U2OS cells were treated with 1 µg/mL doxycycline for 48 hours before fixation and staining with DAPI. Nuclear sizes were analyzed by measuring DAPI area using Photoshop software. The average nuclear area of cells overproducing Cdt1-WT was 1.2 fold larger than control cells, whereas cells expressing Cdt1-5A had even larger average nuclear area (∼1.7 fold higher than control cells). Representative results of two independent experiments are shown; total numbers of cells analyzed is listed under the histograms. Asterisks indicate statistical significance (*** p<0.001, ** p<0.01) determined by Mann–Whitney U-test. Mean +/- standard deviation is indicated.

**B)** U2OS cells were treated as indicated in (**A)** and stained with an anti-γ-H2AX antibody (green). Nuclei were stained with DAPI (blue). Representative results of two independent experiments are shown. Quantification of the percentage of γ-H2AX positive cells is shown with the total number of cells analyzed listed under the histogram.

**Figure S3. Cdt1 is phosphorylated to inhibit DNA re-replication.**

Quantification of the experiments in (Fig. 1B and 1C) showing all cell cycle phase distributions (G1, S, G2/M, and re-replication). n >4.

**Figure S4. Cdt1 mobility by Phos-Tag gel analysis and tests of inhibitor activities.**

**A)** Asynchronously proliferating U2OS cells ectopically expressing HA-tagged Cdt1-WT were treated with 20 J/m^2^ UV 60 minute prior to harvest to induce degradation of Cdt1 (lane 1). Cells were also synchronized in G1 phase by nocodazole arrest and release for 3 hrs (lane 2) or held in nocodazole plus MG132 to induce Cdt1 hyperphosphorylation (lane 3). Lysates of arrested cells were either mock-treated (lane 3) or incubated with lambda and CIP phosphatase (lane 4) for 30 minutes. The samples were then subjected to Phos-tag SDS-PAGE followed by immunoblotting with HA antibody.

**B)** U2OS cells were treated as indicated in Fig. 3C. Mitotic phosphoproteins were analyzed by immunoblotting with an anti-Mpm-2 antibody, a mitotic marker that recognizes a large subset of mitotic phosphoproteins and is sensitive to CDK1 activity in M phase [56].

**C)** U2OS cells were mock treated (lane 1) or treated with 6 µM CVT313 for 6 hours (lane 2), then probed for endogenous Cdc6. Cdc6 is stabilized by CDK2/Cyclin E activity during late G1 phase, and its degradation reflects loss of CDK2-mediated stabilization [57].

**D)** U2OS cells were mock treated (lane 2), treated with 20 J/m^2^ UV (lane 1), or arrested in G2/M phase (lane 3) followed by 30 µM SB203580 treatment (lane 4) for one hour. The mitogen-activated protein kinase-activated protein kinase 2 (MK2) is a direct substrate of p38 [84]. The phosphorylation and total protein levels of MK2 were analyzed by immunoblotting. PonceauS total protein stain serves as a loading control, and representative results of two independent experiments are shown.

**Figure S5. Cdt1 dephosphorylation is inhibited by calyculin A (CalA) and high-dose okadaic acid (OA).** U2OS cells arrested with nocodazole were treated with MG132 and CDK1 inhibitor (lanes 4, 6, and 8) to induce dephosphorylation. As indicated, cells were pre-treated for one hour with okadaic acid (OA, lanes 5-8) or with calyculin A (CalA lanes 3-4) at the indicated concentrations. Okadaic acid inhibits PP2A at low concentrations and can only inhibit PP1 at high concentrations [61]. Cells were harvested by mitotic shake off, and whole cell lysates were subjected to standard SDS-PAGE followed by immunoblotting with HA antibody. A representative of two independent experiments is shown.

