## supplemental figures for Zhou, Pozo, et al. for "Distinct and sequential re-replication barriers ensure precise genome duplication"

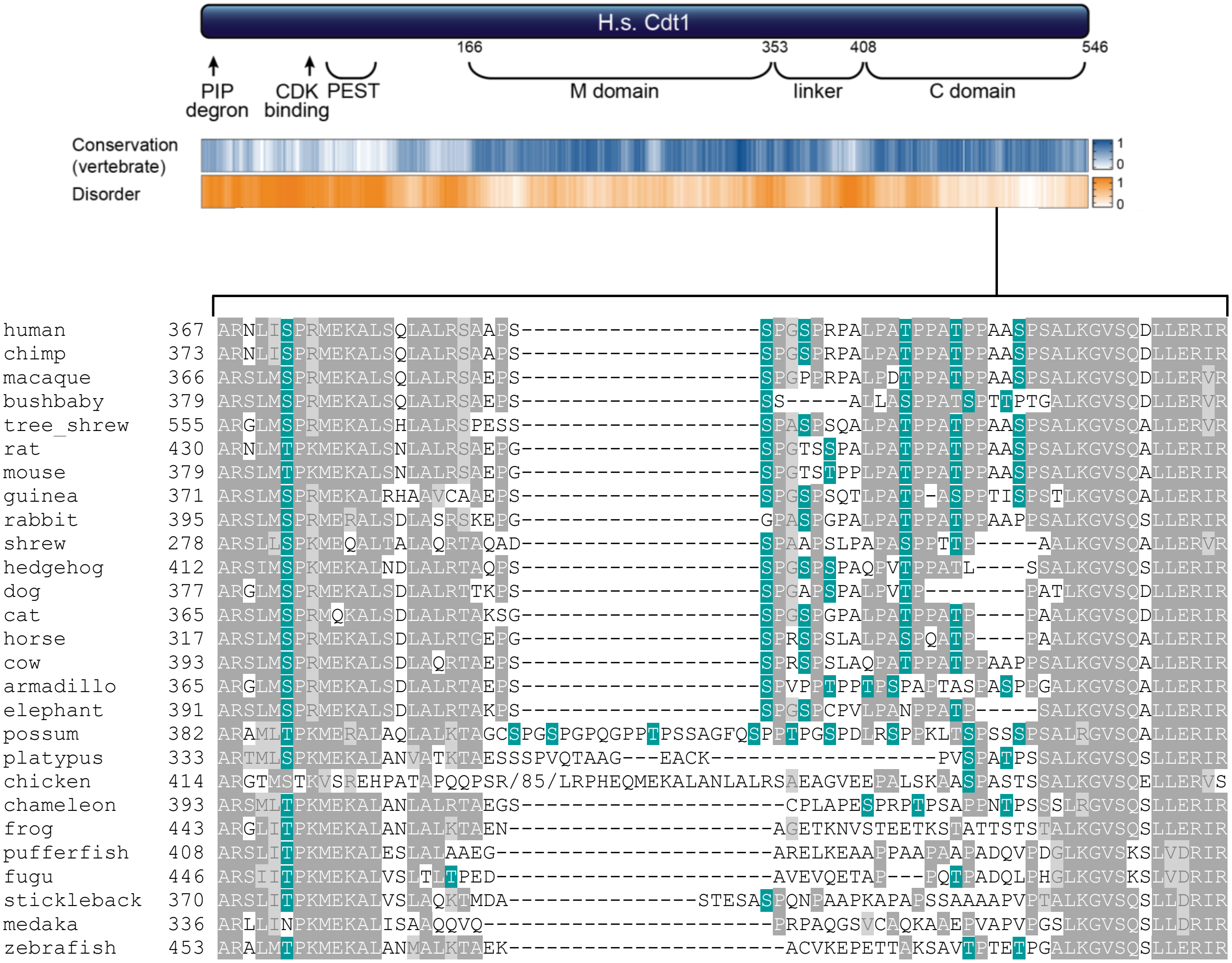


**Figure S1. Cdt1 linker phosphorylation sites in 27 vertebrate sequences.**
A selection of 27 vertebrate sequences for comparison was taken from Miller *et al.* (2007), and Cdt1 protein sequences were retrieved from https://www.uniprot.org/. For the Cdt1 alignment, *Xenopus tropicalis* in Miller *et al.* was replaced with *Xenopus laevis* Cdt1, *Tupaia belangeri* was replaced with *Tupaia chinensis*, and no Cdt1 sequence for *Echinops telfairi* (tenrec) was available. These 27 full-length sequences were aligned with ClustalW at https://www.genome.jp/tools-bin/clustalw using the default settings, and the resulting alignment was visualized with BoxShade, 50% identity or similarity were shaded medium and light grey (https://embnet.vital-it.ch/software/BOX_form.html ). The portion corresponding to the Cdt1 linker domain is shown using common names. All potential CDK/MAPK phosphorylation sites are shaded green, and an 85 residue insertion in chicken Cdt1 lacking any potential CDK/MAPK phosphorylation sites was deleted for clarity. The 27 sequences are from the following species: *Homo sapiens, Pan troglodytes, Macaca mulatta, Otolemur garnettii, Tupaia chinensis, Rattus norvegicus, Mus musculus, Cavia porcellus, Oryctolagus cuniculus, Sorex araneus, Erinaceus europaeus, Canis familiaris, Felis catus, Equus caballus, Bos Taurus, Dasypus novemcinctus, Loxodonta Africana, Monodelphis domestica, Ornithorhynchus anatinus, Gallus gallus, Anolis carolinensis, Xenopus laevis, Tetraodon nigroviridis, Takifugu rubripes, Gasterosteus aculeatus, Oryzias latipes,* and *Danio rerio.*

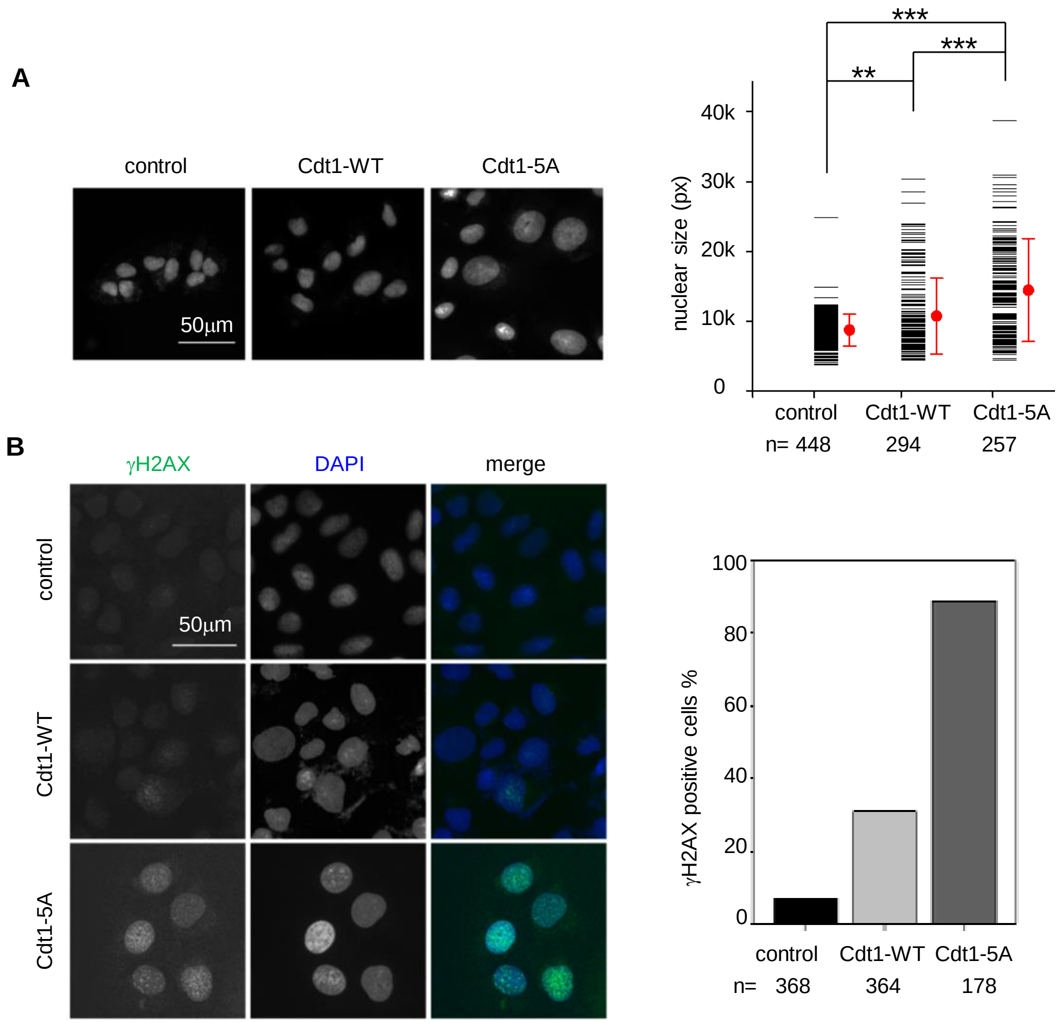


**Supplementary Figure S2. Unphosphorylatable Cdt1 induces giant nuclei formation and DNA damage.**
**A**) U2OS cells were treated with 1 µg/mL doxycycline for 48 hours before fixation and staining with DAPI. Nuclear sizes were analyzed by measuring DAPI area using Photoshop software. The average nuclear area of cells overproducing Cdt1-WT was 1.2 fold larger than control cells, whereas cells expressing Cdt1-5A had even larger average nuclear area (~1.7 fold higher than control cells). Representative results of two independent experiments are shown; total numbers of cells analyzed is listed under the histograms. Asterisks indicate statistical significance (*** $p<0.001$, ** $p<0.01$) determined by Mann–Whitney U -test. Mean +/- standard deviation is indicated.
**B**) U2OS cells were treated as indicated in (**A)** and stained with an anti-γ-H2AX antibody (green). Nuclei were stained with DAPI (blue). Representative results of two independent experiments are shown. Quantification of the percentage of γ-H2AX positive cells is shown with the total number of cells analyzed listed under the histogram.

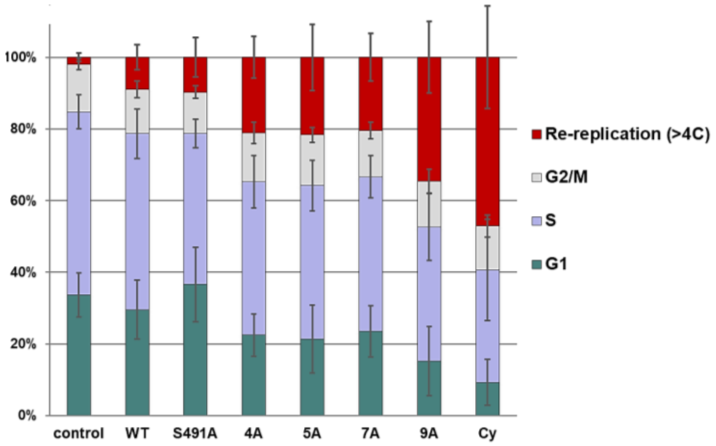


**Supplementary Figure S3. Cdt1 is phosphorylated to inhibit DNA re-replication.**
Quantification of the experiments in (Fig. 1c) showing all cell cycle phase distributions (G1, S, G2/M, and re-replication). n >4.

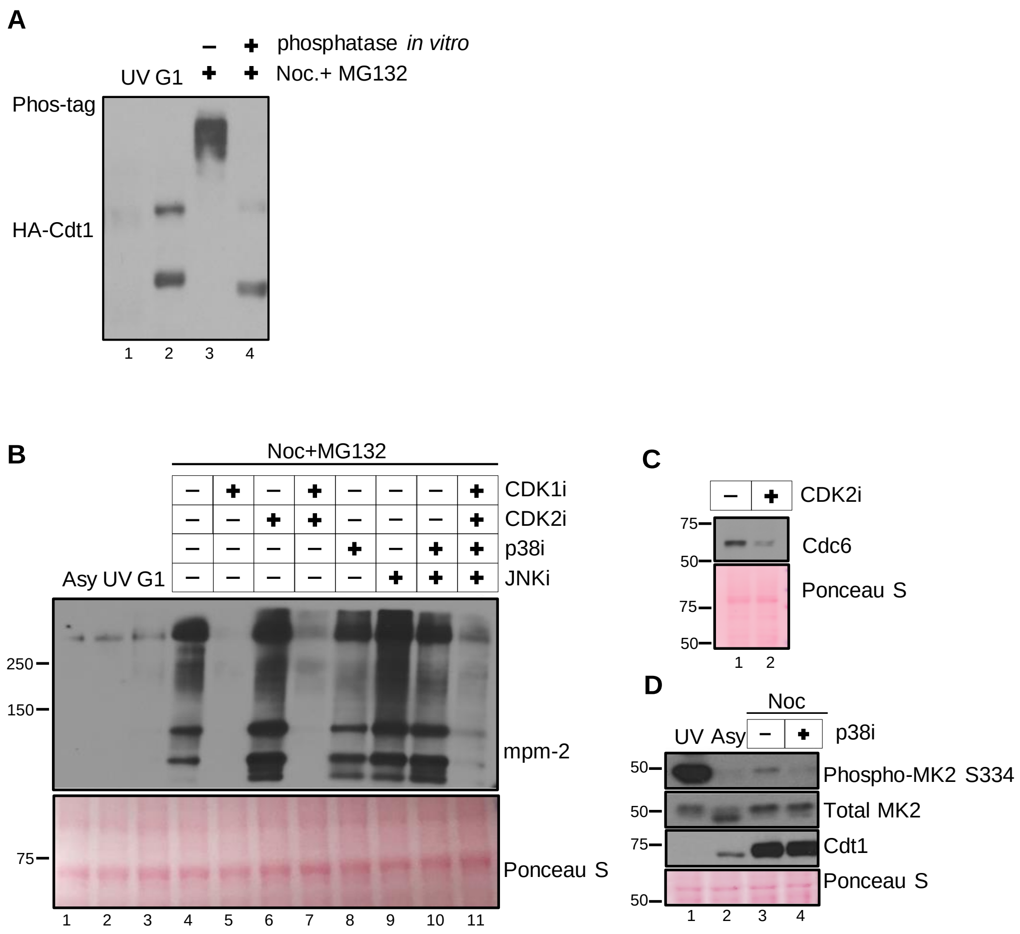


**Figure S4**. **Cdt1 mobility by Phos-Tag gel analysis and tests of inhibitor activities.**

**A**) Asynchronously proliferating U2OS cells ectopically expressing HA-tagged Cdt1-WT were treated with 20 J/m$^2$ UV 60 minute prior to harvest to induce degradation of Cdt1 (lane 1). Cells were also synchronized in G1 phase by nocodazole arrest and release for 3 hrs (lane 2) or held in nocodazole plus MG132 to induce Cdt1 hyperphosphorylation (lane 3). Lysates of arrested cells were either mock-treated (lane 3) or incubated with lambda and CIP phosphatase (lane 4) for 30 minutes. The samples were then subjected to Phos-tag SDS-PAGE followed by immunoblotting with HA antibody.

**B**) U2OS cells were treated as indicated in Fig. 3C. Mitotic phosphoproteins were analyzed by immunoblotting with an anti-Mpm-2 antibody, a mitotic marker that recognizes a large subset of mitotic phosphoproteins and is sensitive to CDK1 activity in M phase (Yaffe *et al.* 1997).

**C**) U2OS cells were mock treated (lane 1) or treated with 6 µM CVT313 for 6 hours (lane 2), then probed for endogenous Cdc6. Cdc6 is stabilized by CDK2/Cyclin E activity during late G1 phase, and its degradation reflects loss of CDK2-mediated stabilization (Mailand *et al.* 2005).

**D**) U2OS cells were mock treated (lane 2), treated with 20 J/m$^2$ UV (lane 1), or arrested in G2/M phase (lane 3) followed by 30 µM SB203580 treatment (lane 4) for one hour. The mitogen-activated protein kinase-activated protein kinase 2 (MK2) is a direct substrate of p38 (Stokoe *et al.* 1992). The phosphorylation and total protein levels of MK2 were analyzed by immunoblotting. PonceauS total protein stain serves as a loading control, and representative results of two independent experiments are shown.

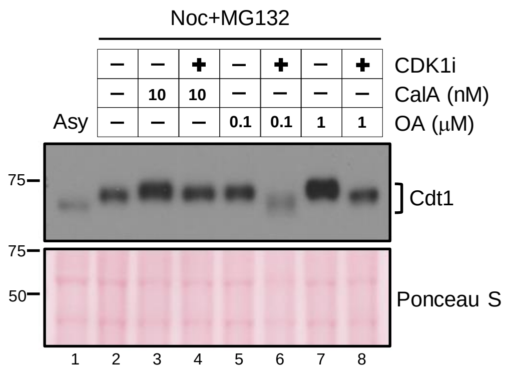


**Supplementary Figure S5. Cdt1 dephosphorylation is sensitive to calyculin A (CalA) and high-dose okadaic acid (OA).** U2OS cells arrested with nocodazole were treated with MG132 and CDK1 inhibitor (lanes 4, 6, and 8) to induce dephosphorylation. As indicated, cells were pre-treated for one hour with okadaic acid (OA, lanes 5-8) or with calyculin A (CalA lanes 3-4) at the indicated concentrations. Okadaic acid inhibits PP2A at low concentrations and can only inhibit PP1 at high concentrations (Swingle *et al* 2007)]. Cells were harvested by mitotic shake off, and whole cell lysates were subjected to standard SDS-PAGE followed by immunoblotting with HA antibody. A representative of two independent experiments is shown.
